# Natural disaster alters the adaptive benefits of sociality in a primate

**DOI:** 10.1101/2023.07.17.549328

**Authors:** C. Testard, C. Shergold, A. Acevedo-Ithier, J. Hart, A. Bernau, JE. Negron-Del Valle, D. Phillips, MM. Watowich, JI. Sanguinetti-Scheck, MJ. Montague, N. Snyder-Mackler, JP. Higham, ML. Platt, LJN. Brent

## Abstract

Weather-related disasters can radically alter ecosystems. When disaster-driven ecological damage persists, the selective pressures exerted on individuals can change, eventually leading to phenotypic adjustments. For group-living animals, social relationships are believed to help individuals cope with environmental challenges and may be a critical mechanism enabling adaptation to ecosystems degraded by disasters. Yet, whether natural disasters alter selective pressures on patterns of social interactions and whether group-living animals can, as a result, adaptively change their social relationships remains untested. Here, we leveraged unique data collected on rhesus macaques from 5 years before to 5 years after a category 4 hurricane, leading to persistent deforestation which exacerbated monkeys’ exposure to intense heat. In response, macaques increased tolerance for and decreased aggression toward other monkeys, facilitating access to scarce shade critical for thermoregulation. Social tolerance predicted individual survival for 5 years after the hurricane, but not before it, revealing a clear shift in the adaptive function of social relationships in this population. We demonstrate that an extreme climatic event altered selection on sociality and triggered substantial and persistent changes in the social structure of a primate species. Our findings unveil the function and adaptive flexibility of social relationships in degraded ecosystems and identify natural disasters as potential evolutionary drivers of sociality.

**One-Sentence Summary:** *Testard et al.* show that a natural disaster altered selection on sociality in group-living primates triggering persistent changes in their social structure.

## MAIN TEXT

With the intensifying climate crisis, extreme weather events are expected to become less predictable and increase in both frequency and force (*1*, *2*). Weather-related natural disasters can cause sustained destruction of the natural landscape, depleting resources, and degrading infrastructure upon which humans and other animals depend (*3*). Persistent disaster-induced ecosystem modifications may then alter the selective pressures exerted on individuals living within them, which can, over time, enforce changes in their traits (*4*). Group-living animals may cope with environmental upheavals by adaptively changing their patterns of social interactions, with potential implications for the transmission of genes, information and disease (*5*, *6*). Whether natural disasters alter selective pressures on patterns of social interactions by changing the costs and benefits individuals gain from them, and whether group-living animals can adaptively modify their social relationships as a result remains untested. The lack of empirical evidence on this topic arises from the difficulty of studying behavioral responses to natural disasters, which are unpredictable and cannot be reproduced in the laboratory, and the rarity of long-term demographic data required to capture fitness consequences of biological traits.

On September 20^th^ 2017, Hurricane Maria made landfall as a Category 4 hurricane on Puerto Rico, causing devastating destruction and a human death toll reaching the thousands, one of the deadliest hurricanes in Caribbean history (*7*). A population of rhesus macaques living on a small island off the coast of Puerto Rico, Cayo Santiago, was also hit by this storm (**Fig. 1A**). Hurricane Maria destroyed 63% of Cayo Santiago’s vegetation, along with nearly all existing research infrastructure (**Fig. 1B-C**). Five years later (Spring 2022), the tree canopy cover remains far below pre-hurricane levels (**Fig. 1B, S1A**), and surface temperatures on the island regularly exceed 40°C (**Fig. 1D, S1C**). Hurricane-induced deforestation resulted in the majority of the island being exposed to these high temperatures. From 2018 to 2021, 48% of sample points from temperature loggers deployed on exposed (i.e., non-shaded) parts of the island detected temperatures above 40**°**C during the day (10am - 5pm) (N=190,240 sample points from 25 temperature loggers, see **Methods**, **Fig. S1C**). Such extreme temperatures compromise the ability to regulate internal temperature, and can result in heat cramps, heat exhaustion, heatstroke, and hyperthermia (*8*, *9*). Primates have limited capacity for physiological thermoregulation (*10*, *11*) and therefore often cope with extreme temperatures by changing their behavior (*12–14*). Cayo Santiago’s macaques address thermoregulatory challenges posed by the hot Caribbean sun primarily by accessing a fundamental resource, shade (on average 7**°**C cooler than exposed areas, **Fig. 1D**, t-test, *P*<0.001).

**Fig. 1.**
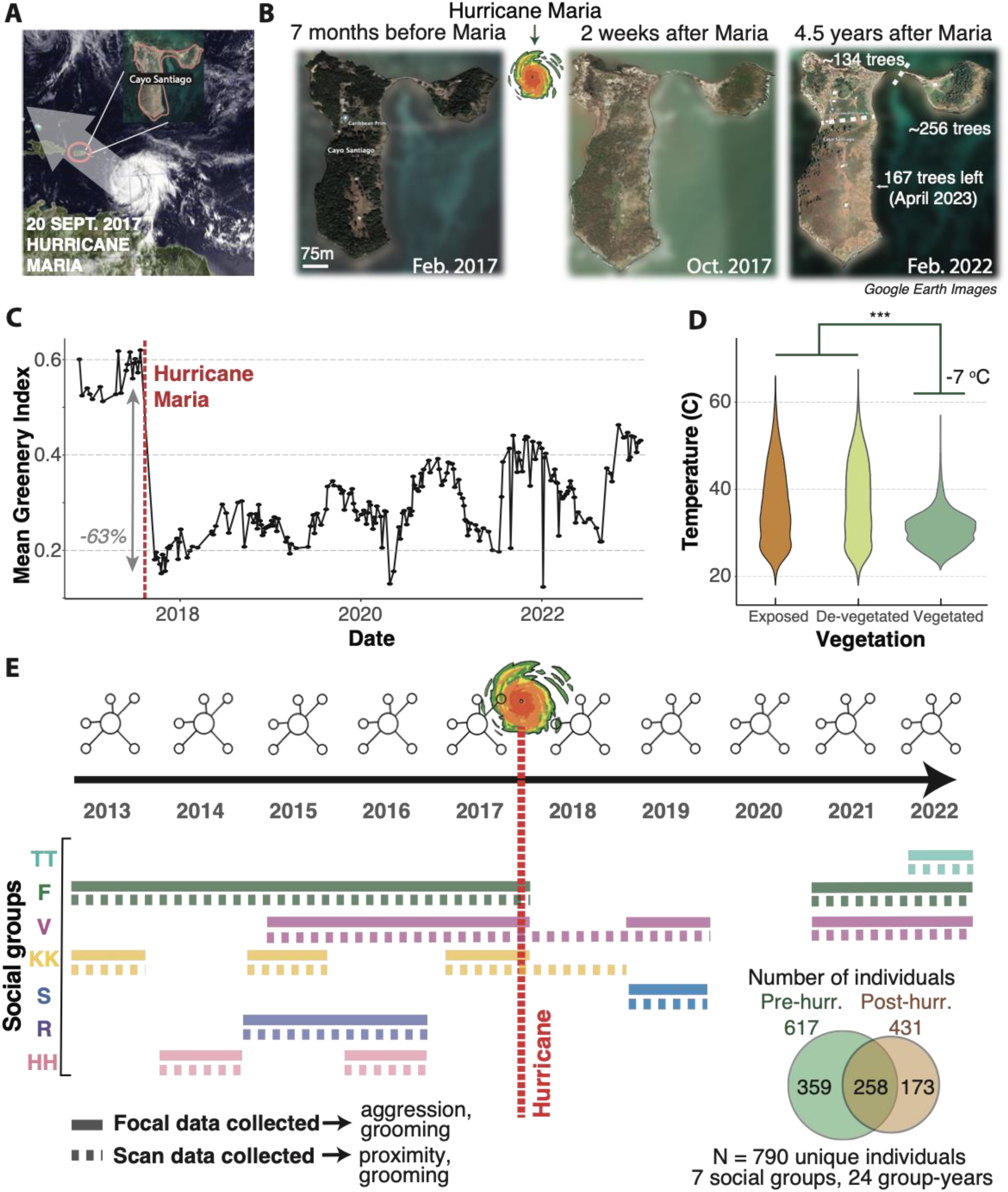
Hurricane Maria caused a persistent drop in vegetation and a concomitant increase in surface temperature. (**A**) Left: Path of Hurricane Maria making landfall on Puerto Rico, including Cayo Santiago, on September 20th 2017. VIIRS image captured by the National Oceanic and Atmospheric Administration’s Suomi NPP satellite. (**B**) Google Earth Images taken at three time points: 7 months before Maria (left), 2 weeks after Maria (middle) and 4.5 years after (right). Tree census conducted in April 2023 revealed only 167 trees were left on the upper part of “Big Cayo” (see **Fig. S1** for satellite images from more time points and tree census on the rest of Cayo Santiago). (**C**) Foliage cover from Cayo Santiago Island, as measured by greenness from satellite imagery (NDVI), decreased by 63% in the 3 months following Hurricane Maria (thick gray bar). Note that NDVI index captures all types of vegetation, including grass and small bushes. Although satellite NDVI increases steadily up until 2022, this largely reflects grass (**Fig. S1**) which does not provide shade. Even when including all vegetation types, vegetation cover still remains far below pre-storm levels. (**D**) Distribution of temperatures post-hurricane (2018–2022) in three types of terrain: *Exposed* - areas without vegetation cover both before and after Hurricane Maria; *De-vegetated* - areas which lost their vegetation cover after Hurricane Maria; and *Vegetated* - areas with vegetation cover. Temperatures in vegetated areas are lower than in other areas (t-test, p<0.001). (**E**) Behavioral data (scans and focal follows) collected from 5 years before Maria to 5 years after, across 7 social groups. Full lines: focal follows; Dotted lines: scans. Inset: Venn diagram of the number of individuals pre- and/or post-hurricane.

Shade thus became a scarce and critical resource after hurricane Maria. Based on evidence showing increased competition when resources are scarce or monopolizable (*15–17*), after the disaster we predicted lower social tolerance and more frequent aggression in rhesus macaques, who are widely regarded as among the most despotic of primate species (*18*). Previously we reported that, instead of becoming less prosocial, macaques on Cayo Santiago dramatically increased their tolerance for other monkeys in the year immediately following Hurricane Maria (*19*). Those initial results led us to hypothesize that increased social tolerance was an adaptive response–monkeys were using their social networks to access now scarce shade to cope with hurricane-induced devegetation and exacerbated heat stress. In the current study we test this hypothesis and predict that changes in social structure detected in the immediate aftermath of the hurricane would persist, and would have direct fitness consequences. We tested these predictions using 10 years of behavioral and demographic data from before and after the disaster (**Fig. 1E**). We found that hurricane Maria altered the adaptive benefits of social relationships in this highly despotic species (*20*) and triggered substantial and persistent changes in the population’s social structure.

### Social tolerance was higher, and aggression was lower up to 5 years after the hurricane

To assess the long-term impacts of Hurricane Maria on social relationships, we used behavioral data collected during a 10-year period from 5 years before Hurricane Maria through 5 years after (pre-hurricane years: 2013-2017; post-hurricane: 2018-2022, excluding 2020 due to the Covid19 pandemic, **Fig. 1E**) in 7 distinct social groups. A total of 790 unique adult individuals with behavioral data before and/or after Hurricane Maria were included in this analysis. We focused our analyses on relationships in which partners spent time in close proximity to each other (i.e., within 2m (*21*)). Whether or not pairs of individuals tolerate being close to each other is an active choice (*22–24*) which is repeatable and indicates an affiliative relationship between them (*25*). For these reasons, proximity is widely used to quantify social structure in primates (*26–29*). Indeed, proximity in close space is tightly linked to resource sharing: macaques that do not wish to share a resource do not tolerate others in close proximity (*23*, *30*, *31*). Reduced aggression between individuals in close proximity further indicates social tolerance, so we also quantified changes in aggression.

We constructed proximity and aggression networks for each group in each year from 2013 to 2022 and compared individuals’ propensity to be in proximity or aggressive to conspecifics before and after the hurricane (details in **Methods**). We used a linear mixed model to compare proximity and aggression in each post-hurricane year to pre-hurricane years (pooled), controlling for the following potential confounding variables: group size, individual age, and sex (see **Methods**). We also conducted a year-by-year analysis taking 2013, the first year in our data, as a baseline for comparing all subsequent years (see **Fig. S3, Methods**). Our models accounted for uncertainty in edge weights due to unequal sampling across individuals, groups and years (see **Methods** (*32*)).

We found that increased social tolerance persisted for 5 years after Hurricane Maria, peaking in the year immediately following the disaster (**Fig. 2A-B**). In the year following Hurricane Maria, Cayo Santiago macaques showed an acute response in which the propensity to be in proximity to another monkey was on average 1.72 standard deviations higher–more than triple–compared to pre-hurricane years (95% CI = [1.60 - 1.83], **Fig. 2C, Data S1**; mean pre-hurricane was 0.18 compared to 0.58 in 2018). Two-to-five years after the storm, proximity amongst macaques remained between 0.47 and 0.84 standard deviations higher than pre-hurricane levels (*P*<0.001 for all years post-hurricane, **Fig. 2C, Data S1**). Similarly, we found a higher number of proximity partners multiple years following the hurricane, peaking immediately after the disaster (**Fig. S2, Data S1**). Population density over the whole island remained stable over the 10 years analyzed and therefore cannot explain these results (i.e, monkeys did not simply interact with others at higher rates because there were a greater number of available partners: mean total population size = 1498 in pre-hurricane years; mean = 1491.2 in post-hurricane years). Given the lack of vegetation recovery in years 2-5 post-hurricane (**Fig. 1B-S1**), the acute component of this social response (i.e. large peak in the year immediately following Maria) indicates a partial decoupling from changes in the environment that are sustained over years following the disaster. This acute component might reflect stronger social cohesion, which is often observed in humans following both natural disasters (*33*) and terrorist attacks (*34*).

**Fig. 2.**
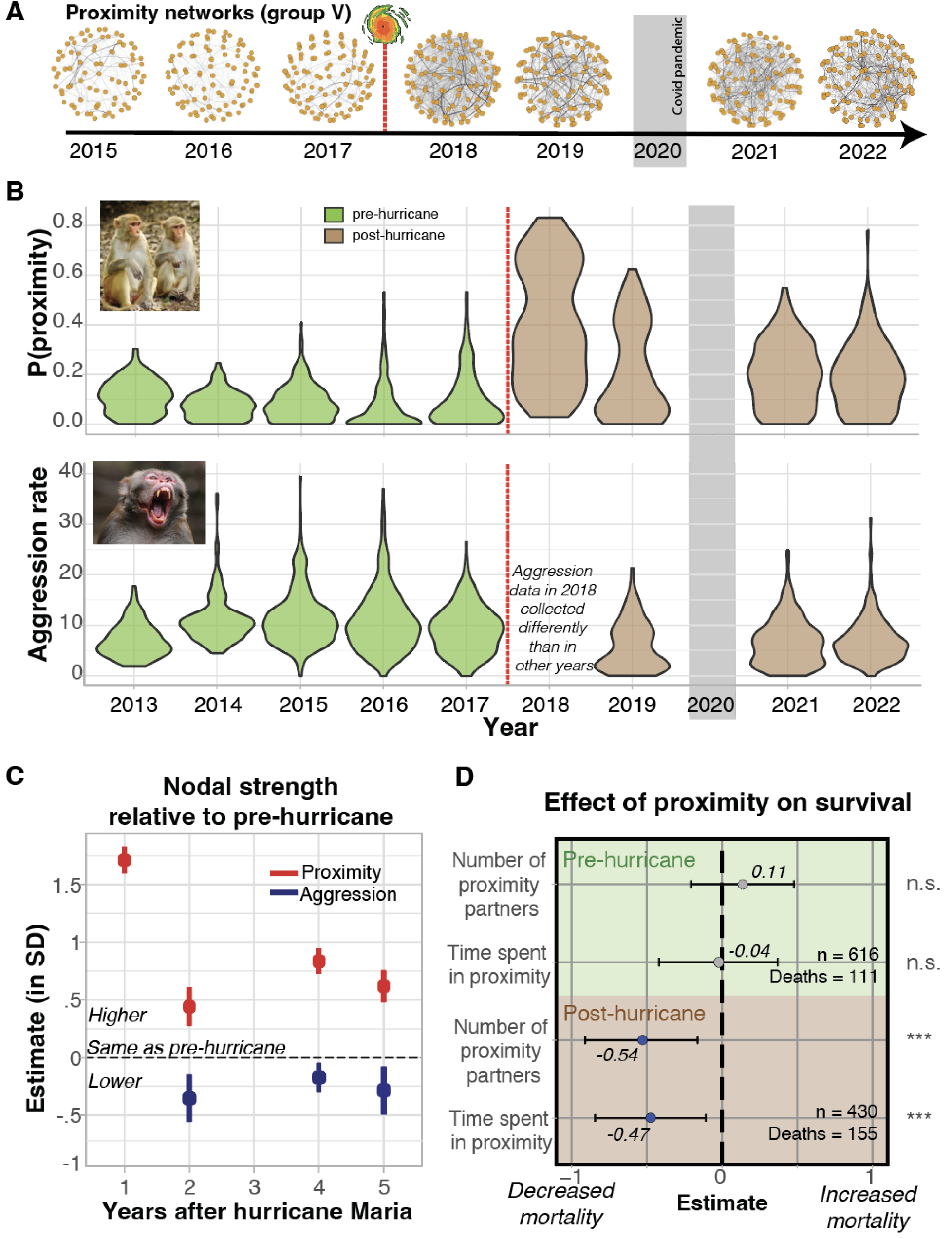
After Hurricane Maria, Cayo Santiago macaques exhibited persistently higher social tolerance, which predicted their survival. (**A**) Proximity networks for social group V from 2013 to 2022, excluding 2020 due to the COVID19 crisis are shown as example networks. Each node represents an individual and edges represent the probability for two individuals to be in proximity to one another. The thicker the line the higher the probability of proximity. Red dotted line marks Hurricane Maria. (**B**) Top: Probability of being observed in proximity to another monkey from 2013 to 2022, across 790 individuals in 6 social groups, 24 group-year combinations. Bottom: same as top panel but for aggression rate (number of aggression events/hours observed). In 2018 aggression data was collected differently such that it was excluded from this visualization (see **Methods**). (**C**) Linear mixed model estimates for the long-term change in standardized proximity and aggression network strength compared to pre-hurricane levels (for grooming results see **Fig. S2**; **Table S1**). Error bars represent the 95% confidence interval (CI), pooled across 100 network iterations (distribution of possible networks given the observed interaction data, see **Methods**). Red: proximity; Blue: Aggression. (**D**) Estimates from survival models for the impact of degree and strength in the proximity networks on mortality risk. Estimates <1: lower mortality risk, >1: higher mortality. Hazard ratio = e^estimate^. Error bars represent the 95% CI pooled across bayesian-generated social networks (see **Methods**). Green: pre-hurricane. Brown: Post-hurricane. ***: p<0.01; n.s.: not significant. Photo credits: Lauren Brent for monkeys in proximity and Gomen S. for aggressive macaque.

Evidence of higher tolerance is corroborated by a sustained reduction in aggression in the years after the disaster, even as resources, particularly heat-reducing shade, were depleted and individuals were more frequently near one another (**Fig. 2B-C**). Lower aggression between members of a despotic species like rhesus macaques in the context of scarce resources and increased proximity was remarkable and unexpected based on published characterisations of this species (*15–17*). Higher tolerance and lower aggression were not associated with consistently higher frequencies of grooming in the five years following the hurricane (**Fig. S2**).

### Persistent environmental degradation changed the adaptive value of social tolerance

Next, we investigated whether higher tolerance predicted survival following Hurricane Maria (study period: September 2017 - August 2022). We tested the effect of two individual-level network metrics from the proximity network: degree (i.e., the number of proximity partners) and strength (i.e., the propensity to be in proximity to others). This analysis included 431 unique adult individuals from 5 social groups, of which 155 died during the study period. Because the values of network metrics changed over the study period, we used time-varying mixed effect cox-proportional hazard models (*35*) to evaluate whether proximity predicted survival from one year to the next during our study period. All models implicitly accounted for age (see **Methods**), sex was included as a fixed effect and social group, individual ID and year as random effects to account for repeated measures, between-group and between-year differences.

We found that both degree and strength of proximity predicted survival following the hurricane, indicating that behavioral responses were adaptive (**Fig. 2D**). Each additional standard deviation in an individual’s number of proximity partners (+12.7 partners) resulted in a 42.69% lower mortality risk (Model estimate = −0.54, Hazard Ratio = 0.58, 95%CI = [0.40 0.86], **Data S1**) and an additional standard deviation in the propensity to be in proximity to another macaque (i.e., proximity strength) led to 37.3% lower mortality risk (Model estimate = −0.48, Hazard Ratio = 0.62, 95%CI = [0.41 0.95], **Data S1**). Focusing on individuals for which we have data before and after the disaster (N_ID_ = 258), we found that inter-individual variation in increased tolerance after the hurricane predicted survival. However, this was only the case for individuals that had low social tolerance before the hurricane (Spearman correlation between plasticity and survival time = 0.84, CI = [0.83 - 0.87], **Fig. S4**, see **Suppl. Text**). This relationship did not hold for individuals that were already highly tolerant pre-hurricane (Spearman’s rho = 0.22, CI = [-0.12 - 0.56]). In other words, being highly tolerant after the disaster is the critical predictor of survival, regardless of how this is achieved (through plastic change or by being highly tolerant to begin with).

To test whether the adaptive benefits of proximity after the hurricane were due to the storm and its aftermath, we tested whether proximity also predicted survival before Hurricane Maria. Using the same modeling approach but focused on *pre*-hurricane years (study period: Jan. 2013 - Sept. 2017; N = 617 individuals, 111 deaths, see **Methods**), we found no evidence that proximity strength or degree predicted survival (**Fig. 2D**, **Data S1**). Furthermore, aggression received or given did not predict survival before or after the hurricane (**Data S1)**. These findings indicate that, by drastically modifying Cayo Santiago’s ecosystem, Hurricane Maria altered the adaptive benefits of tolerating conspecifics in close proximity.

### Stronger thermoregulatory pressure post-hurricane drives the adaptive benefits of social tolerance

The dramatic drop in shade wrought by Hurricane Maria elevated thermoregulatory burdens on Cayo Santigo’s macaques. Temperatures in non-shaded areas of the island regularly exceed 40°C (**Fig. S1C**) and persistent exposure to these temperatures increases the risk of severe heat shock and generally degrades health and physical condition (*8*). Anecdotal reports from observers on the ground suggested macaques used shady areas afforded by the remaining vegetation following Hurricane Maria to lower their body temperature (**Fig. 3A**). From these initial observations, we hypothesized that individuals’ increased proximity to conspecifics after the hurricane and the adaptive benefits of this social response reflected access to scarce shade to deal with the intensified thermoregulatory burden. Since cause of death was not part of our longitudinal data collection protocol, we used temperature, shade status and survival data to test this hypothesis. We found that the temperature difference between shaded and exposed areas was higher in the afternoon (8°C) compared to morning hours (2°C, **Fig. 3B**). Concurrently, macaques were found in areas with more shade in the afternoon (t-test, *P* < 0.01, **Fig. 3C**) and were more likely to be in proximity to a monkey during the hottest hours of the day (Post-hurricane AM mean proximity edge weight = 0.0017; PM = 0.0037, **Fig. 3D**). Finally, we found that proximity during hotter hours of the day (PM) better predicted survival than proximity during cooler daytime hours (AM, **Fig. 3E, Data S1**). These findings endorse the hypothesis that increased tolerance after the hurricane reflects an adaptive behavioral response to increased thermoregulatory stress.

**Fig. 3.**
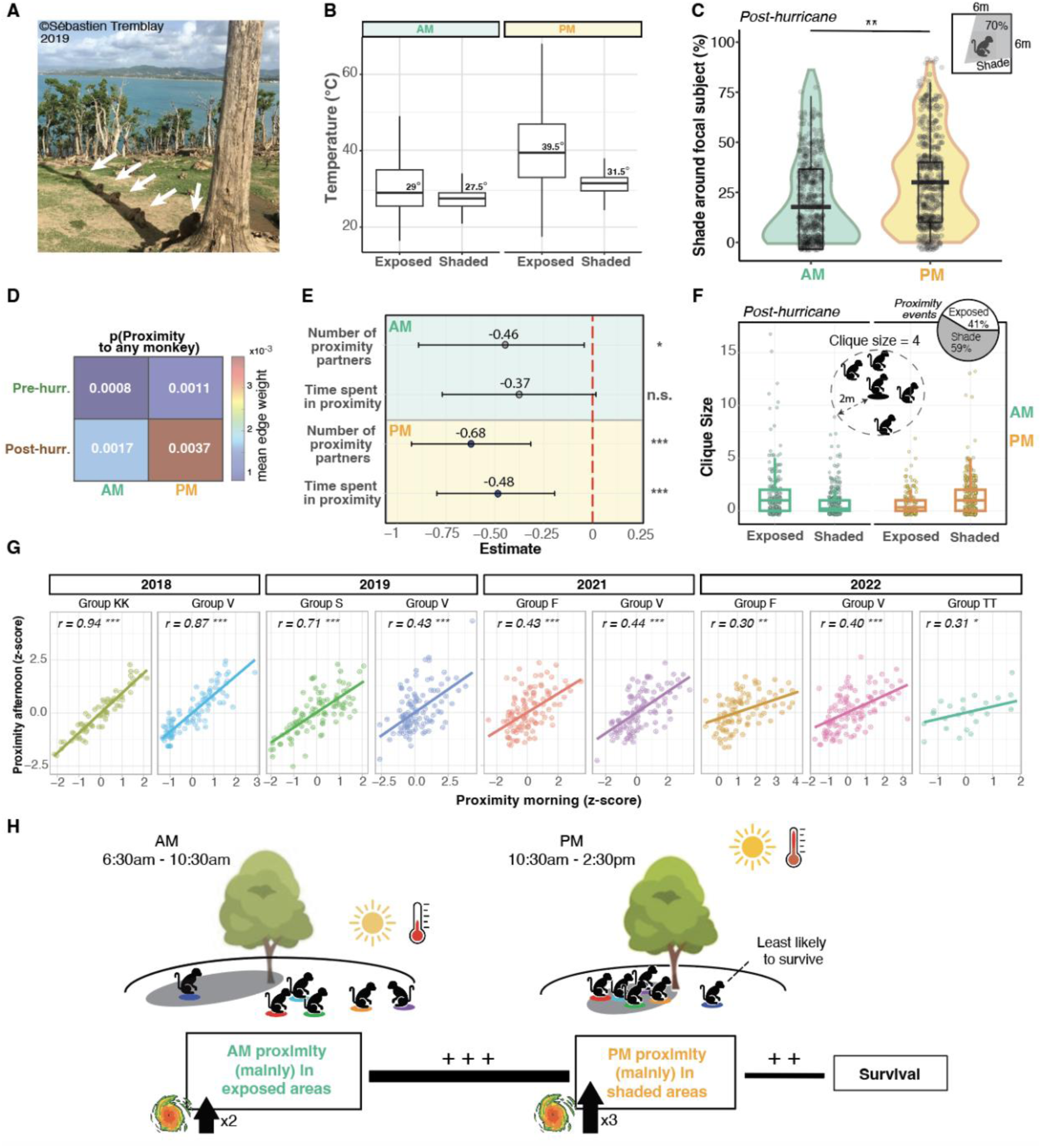
Thermoregulatory needs drive macaques to become more tolerant after Hurricane Maria. (A) Photo taken two years after the storm showcasing Cayo Santiago macaques concentrated in a slim shady area behind a dead tree. (B) Temperature (**°**C) in exposed vs. shaded areas in the morning (left) vs. afternoon (right). (C) Shade (%) in 6m^2^ around focal animal split by morning and afternoon (D) Mean edge weight of proximity networks in the morning vs. afternoon pre- and post-hurricane. (E) Proximity network measures from data collected after the hurricane in the afternoon (orange) better predict survival than from data collected in the morning (green). Bars represent 95% CI. (F) Number of individuals in proximity to focal monkey (i.e., clique size) in exposed vs. shaded areas, morning vs. afternoon. Based on data collected May-August 2019. (G) Spearman’s correlation between morning and afternoon proximity for all groups and years post-hurricane. (H) Schematic diagram summarizing our results: social proximity in the morning doubled after the hurricane compared to before, largely occurs in exposed areas of the island and is strongly correlated with afternoon proximity. Proximity is more likely in the afternoon compared to the morning, mostly occurs in the shade and predicts survival. Thus, the hurricane led to a generalized change in patterns of social interactions, even in non-thermoregulatory contexts, facilitating access to shade when most needed. N.s.: not significant; *: *P*<0.05; **: *P*<0.01; ***: *P*<0.001.

Next, we investigated whether proximity between monkeys after the hurricane solely reflected concentration in cool shaded areas or whether it also occurred in other contexts suggesting a general increase in social tolerance after the hurricane. We collected shade use and proximity data May-August 2019 on 200 individuals across all social groups on Cayo Santiago (see **Methods**). We hypothesized that although social tolerance is most important in the afternoon to access shade, changes in patterns of social interaction will be apparent in other non-thermoregulatory contexts, such as during cool morning hours, reflecting changes in social relationships rather than simply spatial preference for shade during the hottest hours of the day. We found that proximity to conspecifics frequently occurred in exposed areas of the island (41% of observed proximity events), particularly in the morning when temperatures are coolest (**Fig. 3F**). In fact, morning proximity between individuals almost doubled after the hurricane compared to before (**Fig. 3D**). If proximity interactions do reflect social relationships, then we would expect a strong positive correlation between morning and afternoon proximity–which we observe across all groups and years post-hurricane (Spearman correlations, **Fig. 3G**). These results suggest that increased proximity after the hurricane was not simply a byproduct of individuals congregating in narrow shaded spaces during hot hours of the day but also reflects active social decisions in other contexts, which in turn predict access to valuable shade when most needed. In other words, the Cayo Santiago macaques changed how they engage with others and the nature of their social relationships, facilitating access to a scarce thermoregulatory resource in response to intensified heat stress post-disaster (**Fig. 3H)**.

## DISCUSSION

We found that monkeys were persistently more tolerant of others in their vicinity and less aggressive for up to 5 years after Hurricane Maria. Relationships based on social tolerance became more numerous and predicted individual survival after the storm, especially during the hottest hours of the day. These findings support our hypothesis that the adaptive benefits of social tolerance are linked to accessing a thermoregulating resource–shade–in response to increased heat-stress. Monkeys were not simply being passively ‘squeezed’ into now limited shaded spaces but instead showed a generalized increase in social tolerance, including outside of thermoregulatory contexts, suggesting a fundamental change in how they engaged with others. Importantly, social tolerance did not predict survival before the hurricane, demonstrating that hurricane-induced dramatic changes in ecological pressures altered the benefits individuals gain from social relationships, leading to flexible adjustments in who they interact with and how they interact with them. These results show that a natural disaster altered selective pressures on a social phenotype and identify these extreme events as potential evolutionary drivers of sociality in group-living animals.

The disaster-induced change in selective pressures on Cayo Santiago’s macaques and ensuing social flexibility may partially reflect macaques’ inability to leave the island–a situation that is not uncommon amongst animal populations, including humans. Animals living on islands or fragmented habitats (effectively ecological islands) have only limited if not absent opportunities to migrate to new territories when their habitat is degraded (*36–39*). In humans, marginalized populations of low socioeconomic status also suffer from limited capacity to move away from disaster zones (*40*, *41*). Our findings thus generalize to contexts in which natural disasters trap populations in degraded landscapes with limited or absent opportunities to escape, forcing rapid adjustments to new environmental conditions.

Social tolerance following the hurricane was associated with a 42% decrease in mortality risk, an effect comparable to the striking survival impacts of social support in humans (*42*). If fitness-enhancing tolerance is heritable and the environmental conditions favoring this phenotype do not change, hurricane-induced ecological degradation may select for more tolerant individuals, leading to the evolution of an overall more tolerant macaque society on Cayo Santiago in the future. However, the adaptive benefits of increased tolerance may change dynamically over time as new challenges emerge. For example, persistent increases in proximity or close contact may increase disease transmission (*43*), altering the balance between the costs and benefits of social tolerance. The ongoing behavioral and demographic study of this population will provide powerful opportunities to probe the short and long-term impacts of a natural disaster on the evolution of social behavior in a group-living primate.

Our findings demonstrate that rhesus monkeys display considerable social flexibility in response to environmental disruption and that social connections, rather than being static, are dynamically renegotiated to adjust to current demands. The sudden and persistent vegetation drop induced by Hurricane Maria increased competition for shade, which in theory could have provoked higher aggression and greater mortality, but instead elicited greater social tolerance–a remarkable change in social behavior given that rhesus macaques are widely considered to be aggressive and strongly hierarchical (*18*, *20*, *44*). Prior research showed that group-living animals use their social connections to cope with environmental challenges (*45–49*), but whether animals could alter their patterns of social interactions when the environmental challenges they face abruptly change had, until now, remained unclear (*50*, *51*). Our study provides rare evidence of social flexibility in an animal population, which promoted resilience to an unpredictable, extreme, and persistently impactful climatic event. Our study strongly calls into question essentialist assumptions of static species-level behavioral traits (*20*) and shifts focus onto the role of the environment in social behavior. By increasing the frequency of extreme weather events, climate change may ultimately select for species with flexible social systems capable of facultative adjustments to rapidly-changing environments (*52–55*).

## Acknowledgments

We thank the Caribbean Primate Research Center staff for their important roles in data collection and in the maintenance of Cayo Santiago. We also thank Greg Albery, Erin Siracusa, and Delphine de Moor for insightful comments that enriched our analyses and discussion of the results.

## Author contributions

CT and LJNB conceptualized the study; JAH developed methodology specific to this study; CT and CS analyzed the data with input from LJNB and JISS; JN, DP, AA AB and MW collected the data; NSM, JH, MM, MLP, LJNB secured funding for this work; CT and LJNB drafted the manuscript with input from all authors; LJNB supervised this work. All authors read and approved the manuscript.

## Data and materials availability

Upon publication, data will be available on Open Science Framework platform and the code will be on GitHub at https://github.com/camilletestard/Cayo-Maria-Survival

## Supplementary materials

Materials and Methods

Supplementary text

Figs. S1 to S4

Table S1 to S2

References (56–69)

## Supplementary Materials

### Materials and Methods

#### 1. Model system and subject details

We studied a population of rhesus macaques living in a semi free-ranging colony on Cayo Santiago, Puerto Rico (1809 N, 65 44 W). The colony has been almost continuously monitored since it was established in 1938 following the release of 409 animals originally captured in India. Cayo Santiago is managed by the Caribbean Primate Research Center (CPRC), which supplies food to the population daily and water *ad libitum*. Animals are fed once a day in the morning between 7 and 9am. There is no contraceptive use and no medical intervention aside from tetanus inoculation when animals are weaned yearlings. Animals are free to aggregate into social units as they would in the wild. There are no natural predators on the island.

Subjects for this study were adult males and females (at least 6 years old), individually recognizable by tattoos, ear notches, and facial features. We used 7 groups, TT, F, KK, V, R, HH and S, for which there was behavioral data either before or after Hurricane Maria, between 2013 and 2022. Our dataset included 790 unique adult individuals (see **Table S1** for more details per group and year). These groups had home ranges on different parts of the island. Group V ranged on ‘‘small Cayo’’; an area whose land bridge was severed from the main part of island (‘‘big Cayo’’) after the hurricane. In contrast, Group TT, KK, R, HH, S and F ranged on big Cayo with all other groups. We used multiple available years of observational data (HH: 2014, 2016; R: 2015, 2016; F: 2013, 2014, 2015, 2016, 2017; KK: 2013, 2015, 2017, 2018; V: 2015, 2016, 2017) to characterize social behavior before the hurricane (‘‘pre-hurricane’’). Post-hurricane, our sample includes observational data from 2018, 2019, 2021 and 2022 (S: 2019; TT:2022; F:2021,2022; V:2018,2019,2021,2022; KK: 2018). In this study, individuals did not need to have data collected both before and after Maria to be considered.

#### 2. Behavioral data collection

In all years of data collection except 2018 (year immediately following Hurricane Maria) and three social groups in 2016, 2017 and 2019 (HH, KK and S respectively), behavioral data were collected with 10-min focal animal samples, where the duration and partner identity of positive (e.g., grooming) and negative (e.g., aggression, threats, submissions, and displacements) social interactions with adults were recorded. At the 0-, 5-, and 10-min marks of the focal follow, we collected instantaneous scan samples during which we recorded the state behavior of the subject (grooming, feeding, resting, and traveling) and the identity of all adults within two meters (i.e., in proximity). For groups HH in 2016, KK in 2017 and S in 2019, we collected 5-min focal animal samples with instantaneous scan samples at the beginning and end of the focal follow, the data collection protocol was otherwise identical to other non-hurricane years. Importantly for this study, grooming, and proximity were mutually exclusive: grooming took precedence over proximity such that whenever two individuals were grooming they were not recorded as being in proximity as well. We balanced the collection of focal samples on individuals across time of day and across months to account for temporal variation in behaviors. Data-collection hours were from 6am to 2:30pm due to CPRC operating hours. To collect behavioral data before 2020 we used Teklogic Psion WorkAbout Pro handheld computers with Noldus Observer software. After 2020, we used the Animal Observer App on Apple iPads.

In the year following Hurricane Maria (from November 2017 to September 2018), damage resulting in inconsistent access to electricity in Puerto Rico imposed the adoption of a downgraded means of recording data using basic tablets. We recorded group-wide instantaneous scan samples at 10-min intervals. For all animals in view of an observer, we recorded the state behavior of the subject, the identity of their adult social partner when relevant (i.e., if they were grooming) and the identity of all adults within two meters (i.e., in proximity)–similarly to instantaneous scans recorded during focal follows prior to the hurricane. Observers were given 15 mins to complete a group-wide scan session, were required to stand a minimum of 4 m from monkeys and, because of very good visibility of these terrestrial animals, were able to identify them at distances upward of 30 m. While aggressive interactions were recorded during scans in 2018 they were only recorded during focal samples in all other years. Scan samples and focal samples can provide different estimates of brief behaviors like aggression that are not extended in time (*56*). Given this limitation, we excluded 2018 in our quantitative comparison of aggressive rates pre-to-post hurricane in this study.

Our subjects were observed over a mean (SD) of 2.79 (1.87) years per individual, and did not have to be observed both before and after the hurricane to be included in analyses. In other words we did not exclusively conduct within-individual comparisons as in (*19*). Out of 790 subjects, 549 individuals were sampled more than once and 258 (or ∼1/3^rd^) were sampled at least once before and after the hurricane (**Table S2**). In all years but 2018 (where data collection was different), we included on average 4.64 (2.16) h of focal follows and 84.32 (36.96) scan observations embedded within focal follows per individual per year. In 2018, we collected 533.36 (127.41) scans per individual (November 2017 - September 2018). Note that because of the impacts of Hurricane Maria and other storms, data collection stopped on August 31, 2017 and didn’t resume until November 2nd, 2017.

Dominance ranks for individuals were determined separately for each group and year. Rank was also determined separately for males and females. For males, the direction and outcome of win-loss agonistic interactions recorded during focal animal samples or during *ad libitum* observations of a given year was used to determine rank for that year. For females, rank was determined using both outcomes of win-loss agonistic interactions and matriline rank. Female macaques inherit their rank from their mothers, and female ranks are linear and relatively stable over time. In order to account for group size, dominance rank was defined by the percentage of same sex individuals outranked, and ranged between 0 to 100 (0 = lowest rank, outranks 0% of same sex individuals; 100 = highest rank, outranks 100%).

Please note that the following information was not part of the long-term data collection process: whether individuals are in the shade or not and cause of death. Shade status was only collected May-August 2019 (details in following section).

#### 3. Deaths in the aftermath of hurricane Maria

Although the adult death rate peaked in the month after the hurricane (more than triple the expected death rate based on October months in previous years), it returned to expected numbers in subsequent months when compared to previous years (*19*). This peak, however, only represents a small proportion of the population of rhesus macaques (32 individuals died in the month following the hurricane). The lack of mass mortality in the midst of a radically transformed landscape is at least partly explained by the herculean efforts of the CPRC staff to bring food and water to the macaques on Cayo no more than 72 hours after the event.

#### 4. Removal of animals from Cayo Santiago for population management

During the course of the study, the Caribbean Primate Research Center conducted removals of individuals from Cayo Santiago for population control. In 2013, 2014 and 2015 as well as 2021 and 2022, the CRPC removed young individuals (non-adults, <4 yo) from different social groups. As our study focused on adults (> 6yo), removal of young individuals did not impact data collection. In 2016, 2018 and 2019, removals targeted entire social groups (2016: HH, 2018: KK and 2019: S). Removed individuals and groups were chosen based on their genetic profile to optimize the genetic diversity of the Cayo Santiago population. No removals occurred in 2017 due to Hurricane Maria or in 2020 due to the COVID-19 pandemic. A priori selection of individuals to be removed by the CPRC based on social group membership and genetic background excludes the possibility that data collection was biased by removals (e.g. by preferentially removing isolated individuals because they may be easier to catch). In our survival analyses, removed individuals are not considered as ‘dead’, but rather as ‘missing’ (or censored), such that removals did not influence our survival results.

#### 5. Shade use

May-August 2019 we measured shade use of 200 individuals belonging to all groups on Cayo Santiago with a total of 1035 sample points balanced equally between IDs. Each sample point included (1) shade status (Yes or No); (2) Estimated % shade around a 6m radius centered on the focal monkey. 6×6 meters was chosen to give an idea of how much shade is available for the monkeys to use.; (3) The number of individuals of all ages within 2m of the focal monkey (i.e. clique size). If the weather was overcast, how much shade was available was not applicable as this cannot be seen.

#### 6. Changes in vegetation after Hurricane Maria

We quantified changes to vegetation cover as a result of Hurricane Maria, and in particular tree coverage which provides shade, using two quantitative approaches and a qualitative one.

##### a. Greenery index for short-term vegetation drop pre-to-post hurricane

From satellite images, we extracted the ‘‘Normalized Difference Vegetation Index’’ (NDVI) which is the most widely used remote sensing index for assessing vegetation cover (*57*). NDVI is measured using the near-infrared radiation from photosynthetic pigments to assess the photosynthetic activity of vegetation. Importantly, NDVI is sensitive to all types of vegetation, including grass and short bushes which do not provide shade for mid-sized to large vertebrates as tree coverage would. Starting about one year after the hurricane, grass and other ground vegetation started to emerge and are captured by the NDVI index as greenery, even if tree coverage has not recovered (**Fig. S1**). We used NDVI to measure the drop in all vegetation types.

NDVI was extracted from satellite images from Sentinel-hub EO-Browser. We used images from Sentinel 2 L2A, a satellite operated by the US Geological Survey that has a 16-day repeat cycle (i.e., visiting Cayo Santiago every approximately 16 days). Images from Sentinel 2 can be viewed in many formats, including NDVI image format. We created a geojson shapefile of coordinates outlining the entirety of Cayo Santiago, including both the large and small islands, which allowed us to specifically search for images in which there was very little cloud cover over the island. We compiled NDVI scores (0 representing no vegetation and 1 representing full vegetation cover) from satellite images with < 10% cloud cover over Cayo Santiago from January 8th 2017 to August 6th 2018 (approximately 1 year before and after Hurricane Maria). In total, we used 45 images, 19 from before Hurricane Maria (01/08/2017–08/31/2017) and 38 from after (10/26/2017– 08/06/2018). Following Hurricane Maria, vegetation on the island decreased by 63% (*t*-test, *P* < 0.000001).

##### b. Tree census

In March 2023, we undertook a tree census of Cayo Santiago, splitting the island into three areas: Upper Cayo, Mangroves and Small Cayo (see **Fig. S1C**). We censused trees that were ≥ 20cm in diameter at breast height and with leaves covering a minimum of 20% of their branches in an attempt to count only trees that might provide substantial shade (*58*). On Upper Cayo, we performed a total tree count due to the small number of trees remaining and found a total of 167 trees. In the Mangroves and Small Cayo, where tree density was greater, we conducted a random sampling of tree coverage using a circular plot approach. In brief, we counted all trees of the characteristics described above in randomly selected circular plots of 5m^2^ radius. From a starting point, observers walked 30 meters in a compass direction determined via random number generator. That point was the center of the circle of 5m^2^ radius. This process was repeated, without repeated sampling of areas, until approximately 20% of the total area was covered. In Sentinel Hub EO browser we used the area measuring tool to quantify the total area of Small Cayo and the Mangroves, which were 20,000m^2^ and 30,000m^2^ respectively. On Small Cayo, we found a median of 1 tree per 78m^2^. Extrapolating to the total area of Small Cayo (excluding cliff areas) we estimated the total tree coverage to be 20,000/78×1 = 256. For Mangroves, we found a median of 0 trees per 78m^2^ which, using the same approach, yields 0 trees. If using the mean instead of median, we estimated a total of 30,000/78*0.7 = 269 trees in the Mangroves. Upon visual inspection of the google Earth images, the complete tree count likely lies between these two estimates. We did not conduct tree censuses before hurricane Maria such that we were not able to exactly measure the change in tree coverage pre-to-post hurricane.

##### c. Qualitative assessment of Google Earth images

Google Earth images allowed us to visualize the state of Cayo Santiago island at multiple time points from September 2016 until February 2022 (**Fig. S1A**). Although the quality and viewpoints of the images prevented us from performing a quantitative assessment of the tree coverage, they were sufficient for a qualitative assessment and provided valuable historical information about tree coverage pre-hurricane that we do not otherwise have access to. We concluded from this timeline of images and on-the-ground experience that the vegetation cover is far from pre-hurricane levels.

#### 7. Temperature logging after Maria

Surface temperature of the island was quantified using 39 temperature loggers (IButton DS1922L Thermochron Data Logger) dispatched across Cayo Santiago in 2018, i.e., after Hurricane Maria. Loggers were dispatched in three types of terrain: exposed, de-vegetated and vegetated. 12 temperature loggers were located in ‘exposed’ terrain, which did not have tree coverage either before or after the hurricane. 13 loggers were located in ‘de-vegetated’ terrain which used to have tree coverage but did not anymore after the hurricane. Finally, 14 loggers were located in ‘vegetated’ terrain with tree coverage (i.e., in the shade). We compared temperatures from all 39 loggers in the three types of terrain from July 7th 2018 to October 10th 2021, only including temperatures from 6am to 5pm, amounting to a total of 463,510 samples. We find a mean difference of 7°C between shaded areas (‘vegetated’ terrain) and non-shaded areas (exposed and de-vegetated, *P* < 0.00001).

When plotting the mean temperature detected across all loggers in non-shaded areas (i.e., ‘exposed’ or ‘de-vegetated’) by time of day (6am-5pm, considering all months of the year), we find that temperatures exceed 38°C, dangerously intense heat for macaques (*8*), beginning at 10am (**Fig. S1D**). To assess the effect of temperature on sociality and survival patterns (see below for more details), we split our data into cooler morning hours, from 6-10am, and hotter afternoon hours, 10am-2:30pm.

#### 8. Estimating social networks and associated metrics using a Bayesian approach (BISoN)

For each group and year separately, we estimated three types of social networks based on three different social interactions: proximity, aggression and grooming. For each social interaction type, we quantified two individual-level network metrics that allowed us to measure individuals’ sociality: (1) the number of social partners an individual interacted with, or number of connections in the network (i.e., degree); and (2) the propensity at which individuals interacted with their partners, or the sum of probabilities of interaction with all individuals in the network (i.e., strength)(*48*).

Networks are usually constructed by taking a normalized measure of sociality, such as the proportion of sampling periods each pair spends engaged in a social behavior (e.g. the Simple Ratio Index; SRI). These normalized measures ensure that pairs that are observed for longer are not erroneously determined to be more social (*59*). This is important because observing social events can be challenging and uniform sampling over all pairs is not always possible (*60*). Though normalized measures of sociality will not be biased by sampling time, they will be accompanied by varying levels of certainty. For example, edge weights will be treated as a certain 0.5 for both a case where individuals have been seen together once and apart once, and equally where individuals have been together 100 times and apart 100 times. Despite there being considerably more certainty around the values of the latter example than the former, methods to estimate and propagate uncertainty through social network analyses remain largely unused.

In this study, we generated weighted Bayesian networks using the BISoN framework and bisonR package (*32*). This framework allowed us to account for uncertainty in the edges connecting individuals in the network based on how often they were sampled and, more importantly, propagate this uncertainty to subsequent analyses. In the next paragraphs we describe how we modeled each network type in BISoN:

Proximity networks were estimated based on data collected through focal scan sampling (*56*) across all groups and years. Edges in the networks represented the number of times a pair of individuals were observed in proximity relative to the total observation effort for the dyad (i.e., total scans individual A + total scans individual B), or the probability of interaction. Given that the observed interaction data was binary (i.e. for every observation sample, was the focal animal in proximity to another monkey? Y/N) we modeled the uncertainty around the edges using a weak prior, a beta distribution with alpha = 0.1 and beta = 0.1. Proximity networks were undirected.

Aggressive networks, on the other hand, were estimated based on data collected through continuous focal sampling across all groups and years except the year immediately following Hurricane Maria (2018). In 2018 aggressive interactions were collected through scan sampling which makes this year hardly comparable to all other years. As such, 2018 were excluded from our analyses for aggression data. Edges in the network represented the number of aggressive interactions between members of a dyad over the total observation effort for the dyad (i.e., total number of hours observed individual A + total number of hours observed individual B), or the rate of interaction. Given that the interaction data consisted of counts of aggressive events over total time observed, we modeled the uncertainty around the edges using a gamma distribution with alpha = 0.1 and beta = 0.1 as a weak prior. Aggression networks were directed.

Grooming interactions were captured both through scan sampling and continuous focal sampling such that we quantified grooming networks using both approaches described above. Scan-based grooming networks were modeled with a beta distribution (as for proximity) and continuous sampling-based grooming networks were modeled with a gamma distribution (as for aggression). We did not collect continuous focal sampling in 2018 therefore continuous sampling-based grooming networks do not include 2018. Grooming networks were directed.

The BISoN Bayesian framework computes a posterior distribution of possible networks from the observed data, from which we sampled 100 grooming, proximity and aggression networks for each group and year. These networks sampled from posterior distributions were used for downstream analyses. In addition to accounting for uncertainty in edge weights, the BISoN approach benefits from making very few assumptions. The only assumption is that probability/rate is a good characterisation of the edge between two individuals. If edge-weights are based on probabilities of binary events, then binomial is the canonical distribution describing this process. If edge-weights are based on frequency of count events, then Poisson is the canonical distribution describing this process.

Networks generated with BISoN include edges between all dyads by default, as it assumes non-zero probability for all potential interactions, even if that probability is exceedingly small. To compute degree (or number of connections in the network for each individual), we therefore defined a threshold of 0.0001, 1 interaction in 10,000 observations, to differentiate dyads that did interact versus those that did not. After this thresholding, we computed two individual-level network metrics for each possible BISoN network, which were used for downstream analyses: (1) nodal degree using the ‘degree’ function from *igraph* (i.e., number of edges for each individual or node in the network); and (2) relationship strength to all partners in the network using “strength” function in *igraph* (summing all the weights of an individual’s edges in the network). Thresholds of 0.001 or 0.00001 for defining an edge in networks for degree calculations yielded very similar results.

#### 9. Methodological improvements since previous work

Note that we used different analytical methods to conduct network analyses in this study compared to our previous work (*19*). Critically, methods differ in how they account for uncertainty around edge weight estimates, especially for edge weight = 0. In our previous work, we used a subsampling bootstrap approach to control for observation effort differences pre- vs. post-hurricane. However, if no interaction was observed between two nodes in the network, the edge weight would always stay equal to 0 (i.e., no interaction can ever be picked up by sub-sampling) - biasing networks with lower sampling effort toward appearing sparser (with less connections) compared to better-sampled networks, even if observation effort is kept the same through sub-sampling. In this current study, to address this issue, which incidentally pervades the field of animal social network analysis, we used a new Bayesian network method, BISoN, which adds uncertainty around edge weights based on observation effort, such that 0 observed interactions can be modeled as more or less likely to be 0 depending on the observation effort. In pre-hurricane networks, which were based on less sampling effort compared to 2018 ones (∼5-10 times lower), this new method avoids 0-inflation in edge-weights due to sparse sampling. The methodological improvement in this study explains discrepancies with our previous results regarding changes in grooming networks, which were previously reported to be denser in 2018 compared to before the hurricane, but which we now found to be equally dense or sparser in 2018 compared to pre-hurricane years (**Fig. S2**). Note that grooming results differ if we use focal sampling or scan sampling to collect our behavioral data (**Fig. S2**), and as a result patterns of grooming post-hurricane compared to pre-hurricane remain unclear.

#### 10. Long-term change in sociality

To quantitatively assess long-term changes in the propensity to be in proximity pre- to post-hurricane, we used linear mixed models using the R package lme4 (*61*). We included all individuals for which we had behavioral data in any year from 2013 to 2022 across 7 social groups, yielding a total of 790 unique individuals. In all our models our response variable was standardized individual network ‘strength’ (i.e., the sum of weights of an individual’s connections in the network) and our main predictor of interest was the year post-hurricane compared to pre-hurricane years. We included this ‘year post-hurricane’ predictor as a factor in our model, where all pre-hurricane years were combined into a “pre-hurricane” category and post-hurricane years were split into 2018, 2019, 2021 and 2022 (we did not collect data in 2020 due to the Covid19 crisis). We also included covariates age and sex as fixed effects, and social group and individual ID as random intercepts to account for repeated measures and group differences. The same analysis pipeline was also done for grooming and aggression data, as well as individual network ‘degree’ (i.e. the number of unique partners) for all three behaviors.

Finally, to compare changes in proximity in the cooler hours of the day vs. the hotter hours of the day (after 10:30am when temperatures reached 40°C, **Fig. S1**), we repeated the same analytical approach but simply evaluated the effect of hurricane status (pre- or post) and considered data collected earlier in the day (7-10:30am) vs later in the day (10:30am-2:30pm) to construct social networks.

For all models we ran the following assumption checks to ensure model estimates were accurately estimated using the function ‘check_model’ from R package *performance*(*62*): posterior predictive checks, homogeneity of variance, colinearity of fixed effects, normality of residuals, normality of random effects. We did not find violations of the assumptions checked.

#### 11. Mixed effect time-varying Cox-proportional hazard survival models

We investigated whether social tolerance, as measured by the propensity to be in proximity to other monkeys and the number of proximity partners, predicted survival in the years before (2013–2017) and after the hurricane (2018-2022, excluding 2020). We used time-varying mixed-effect Cox proportional hazard models, through the *coxme* R package(*63*), divvying up our analysis by year pre- and post-hurricane. This is because the value of social covariates changed from year-to-year. We used behavioral data from each year to predict survival to the next year. Since we did not collect behavioral data in 2020, this year was excluded from our survival analysis.

For all models, the response variable was a ‘Surv’ object with age in days at the start of the study period, age in days at the time of event (which could be death, removal or end of study), and the event outcome (0: the animal is censored, 1: the animal is dead). Removed individuals during the study period were censored to their removal date and all animals still alive at the end of the study period were censored to the end date. With this approach, removed individuals were not considered as ‘dead’ but rather as ‘missing’ (a common issue in clinical trials(*64*)) and age was accounted for implicitly in all models.

Importantly, study start and end dates varied with the follow-up period considered. In explicit terms: when measuring the survival effects of proximity in 2013, we only considered individuals for which we had behavioral data in 2013. The start of the study was January 1st 2013 and the study end date was December 31st 2013. Individuals who died after 2013 were considered alive when quantifying the effect of 2013 proximity on survival (i.e., event outcome = 0, the animal is censored). All other years were treated the same except 2017 and 2018. For 2017, study end date was set to Sept 20th to exclude post-hurricane deaths. For 2018, study start date was set to Sept 21st to include deaths immediately following Maria. This means that individuals who died immediately of the hurricane were excluded from our analysis (since we do not have behavioral data for them for the Oct. 2017-December 2018 observation year). Overall, we found that out of 790 unique individuals, 111 before the hurricane and 155 died after (total = 266).

We ran separate survival models for pre-hurricane (2013–2017) and post-hurricane (2018-2022, excluding 2020) periods, with either nodal strength or nodal degree in proximity network as social covariates (i.e., a total of four models). All four models also included sex and social status as fixed effects and individual ID, social group, and year considered (or follow-up period) as random effects to account for repeated measures, between-group and between-year differences. All adult monkeys with behavioral data in any year from 2013 to 2022 were considered for this analysis. Pre-hurricane, we considered a total of 616 unique individuals across 5 social groups (HH, R, F, V and KK), of which 111 died in the study period. Post-hurricane, we considered 430 unique individuals across 4 social groups (S, KK, V and F) of which 155 died.

To compare how proximity predicted individuals’ survival post-hurricane earlier in the day compared to later in the day (when hotter), we repeated the same analysis but only considering proximity data collected from 7-10:30am or from 10:30am-2:30pm (i.e., total of four models).

Finally, we repeated the same survival models (i.e. same covariates and random effects) for aggression nodal in-strength and out-strength, running separate models for data collected before and after the hurricane (i.e. total of four models).

For all Cox-proportional hazard models, continuous variables were z-scored. We also ran the following assumption checks to ensure model estimates were accurately estimated: proportional hazard over time using the R function ‘cox.zph’(*65*) and (2) linear relationship between the log hazard and each covariate using R function ‘residuals’.

#### 12. Pooling model results across sampled network metrics

In order to propagate the uncertainty around social networks constructed from the observed interaction data to downstream analyses, all models described above were computed across 100 possible social networks for each group-year-behavior combination generated in BISoN (i.e., extracted from the networks’ posterior distribution). We used the function ‘pool’ from the R package *mice* to combine estimates and the uncertainty around those estimates from the 100 repeated models. As an example, to compute the effect of pre-hurricane proximity on survival, we ran 100 survival models using network metrics extracted from 100 different possible proximity networks per group-year and pooled estimated across those 100 models. We also ran some of these models (testing the effect of proximity degree and strength on survival post-hurricane) using only the observed values, i.e. not pooling results across imputations of network metrics, and found very similar results (**Data S1**).

### Supplementary Text

#### 1. Changes in feeding patterns after Maria do not explain survival results

Before the hurricane, 33.6% of Cayo macaques’ diet was vegetation-based, the rest being ‘monkey chow’ supplemented by the Caribbean primate research center (CPRC) in centralized feeders. Hurricane-induced vegetation loss led to vegetation-based diet dropping to 21.5%, the rest being CPRC-distributed monkey chow. Moreover, starting in April 2017 (6 months before Maria), the CPRC began distributing food across the island resulting in *less* centralized feeding. These feeding changes are unlikely to explain both higher social tolerance post-hurricane and its effect on survival for the following reasons. First, being in proximity to conspecifics did not correlate with the amount of time an individual spent feeding post-hurricane (r=0.06, *P* = 0.10) suggesting one does not enable the other. Second, feeding only occurred once a day in the morning, and afternoon proximity was both highest compared to pre-hurricane and better predicted survival. Reduced aggression post-hurricane, however, may be partly explained by less centralized feeding post-hurricane.

#### 2. Within-individual changes in tolerance from random regression models (i.e., reaction norms) & relationship with survival

To extract pre-to-post rates of change in tolerance for the 258 individuals we have data for before and after the hurricane, we used a mixed effect regression model (“lmer” function in lme4 (*66*)). The response variable was individual proximity strength (z-scored) and fixed effects were hurricane status (pre vs. post), sex and age. We included a random intercept for group and individual ID to model between-group and between-individual variation. We also included a random slope for hurricane status by ID to model within-individual variation in behavior across environments (i,e, before vs. after the hurricane). This random effect can be interpreted as an individual by environment interaction (*67*). The steepness of tolerance changes for each individual pre-to-post hurricane can be calculated by adding up the main fixed effect of hurricane status, the random intercepts for ID and the random slopes for ID. We followed the tutorial from Housley & Wilson 2017(*68*) to conduct this analysis and plot **Fig S4A**.

To determine if reaction norms, i.e. the ability of macaques to become more tolerant after the hurricane, predicted survival, we ran a Bayesian bivariate mixed effect model using brms(*69*) where proximity and survival are both response variables. This modeling approach allows to relate variation in reaction norms with variation in survival outcomes (analysis adapted from tutorial by Tom Houslay “Avoiding the misuse of BLUP in behavioral ecology: II. multivariate modeling for individual plasticity” (*68*)). Plasticity may only be important for individuals with low tolerance levels before the hurricane (i.e. if your tolerance level is already high before the event, it may not be advantageous, or indeed may be impossible, to become even more tolerant afterwards). To account for baseline tolerance, we split our population in two and ran two separate bivariate models: one for low tolerance individuals pre-hurricane (less than average) and one for high tolerance individuals pre-hurricane (more than average).

Proximity was z-scored and modeled with a gaussian distribution. Hurricane status, sex and age were included as fixed effects. We also included random intercepts for group and ID and random slope for hurricane status by ID. Survival time (number of days in the study post-hurricane) was right censored (0= alive or removed; 1:died in the years following the hurricane). Removed individuals were not considered as dead. We used a Weibull distribution to model this censored time variable. Sex and age at the time of event (death, removal or end of study) were included as fixed effects. We also included random intercepts for group and id. We found that inter-individual variation in plasticity predicted survival only for individuals which had low social tolerance before the hurricane (correlation between reaction norms and between-individual variation in survival time = 0.84, CI = [0.83 - 0.87], **Fig. S4B**). This relationship did not hold for individuals which were already highly tolerant pre-hurricane (rho = 0.22, CI = [-0.12 - 0.56]).

### Supplementary Figures

**Fig. S1.**
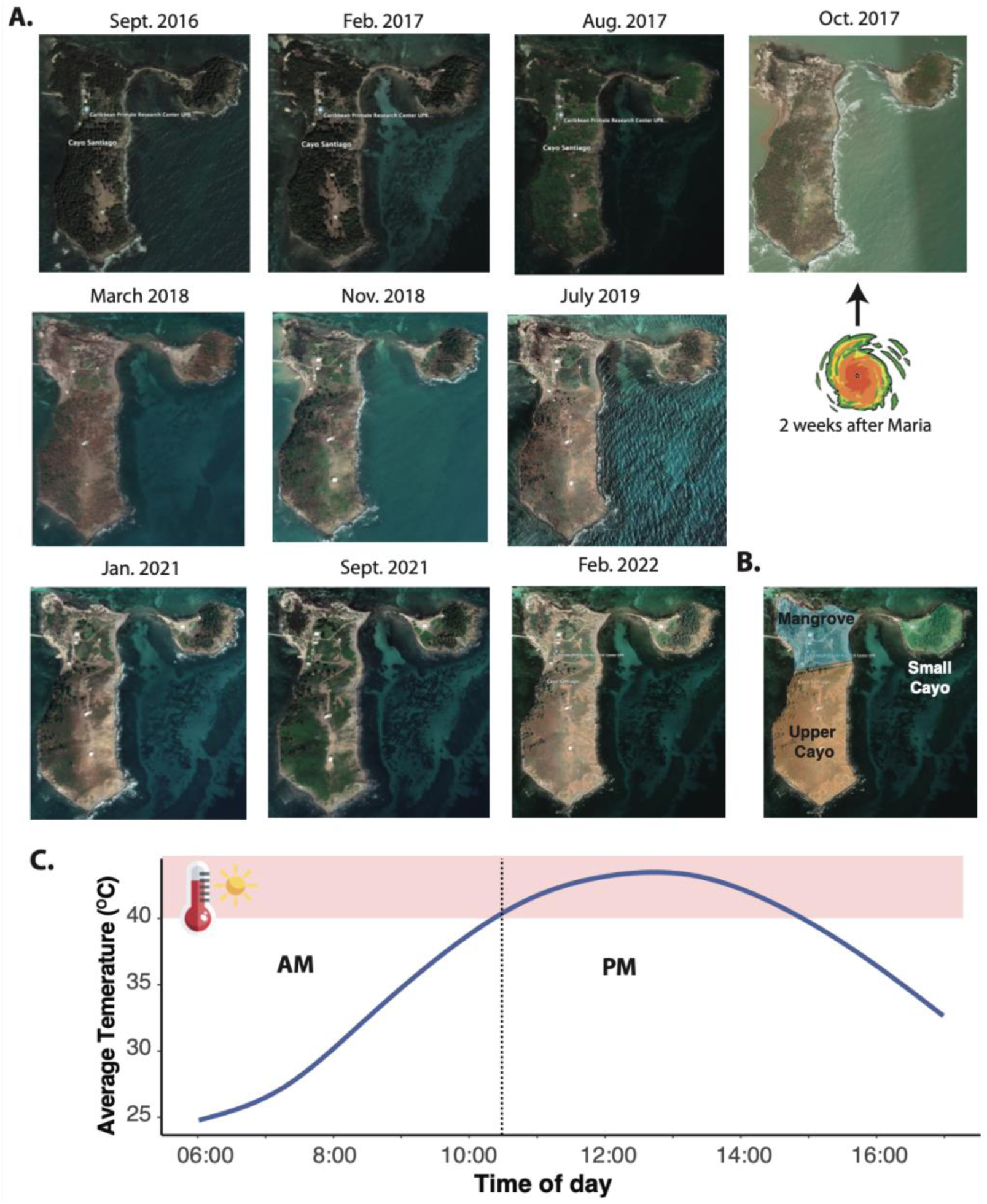
Vegetation and temperature after Hurricane Maria. (A) Timeline of Google Earth Satellite images of Cayo Santiago. (B) Map of the sections of Cayo Santiago for tree census. (C) Average day temperatures from 2018 to 2023 in exposed temperature loggers (average over N=190,240 sample points from 25 temperature loggers, 6am-5pm). Dotted line indicates the AM/PM divide.

**Fig. S2.**
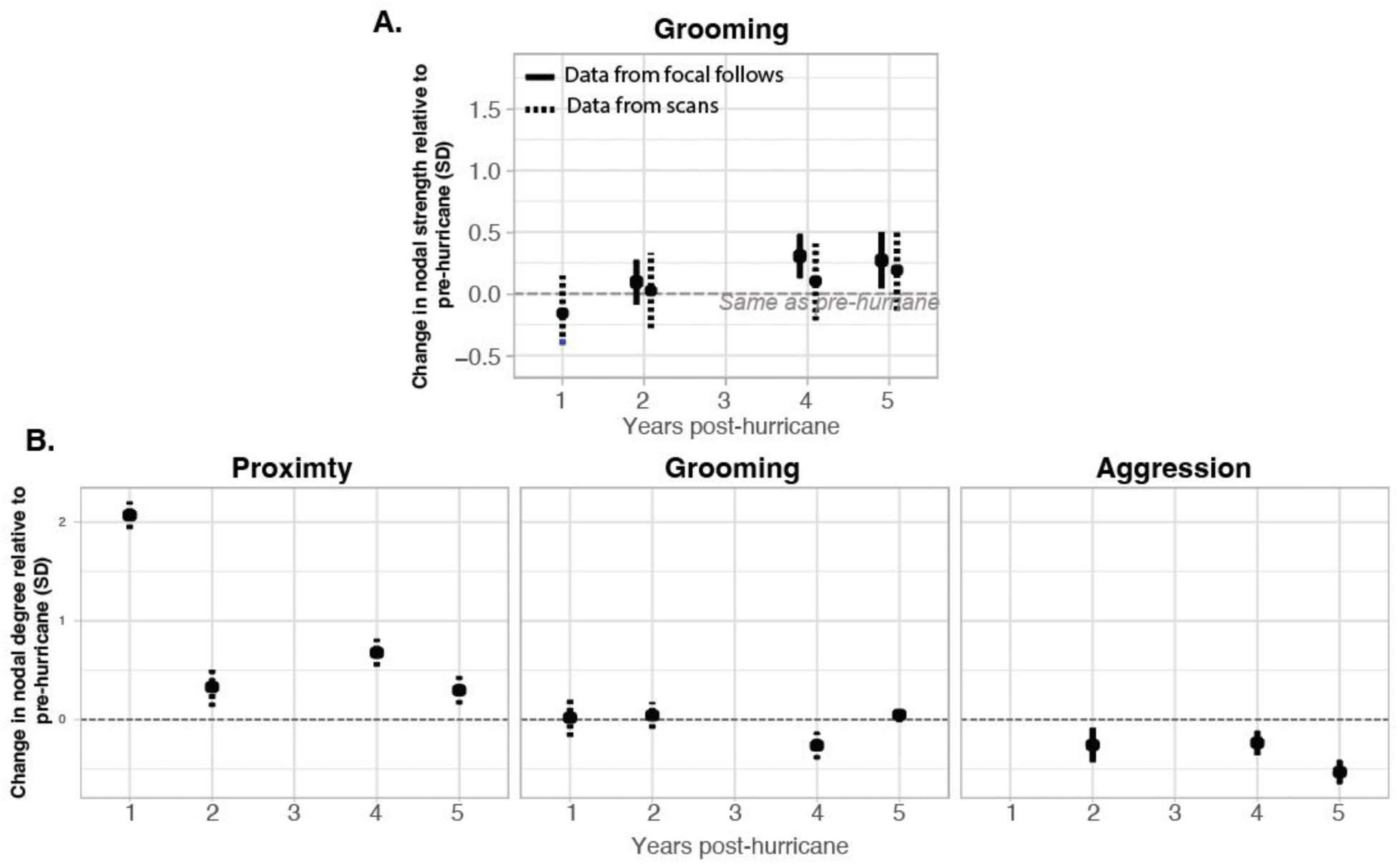
(A) Long term effect of Maria on grooming network strength. Results for linear mixed models comparing 2018-2022 to pre-hurricane years. (B) Long term effect of Maria on network degree for proximity, grooming and aggression. Results for linear mixed models comparing 2018-2022 to pre-hurricane years. Dotted line represents model outputs from data collected with scan samples. Full line from data collected with focal follows.

**Fig. S3.**
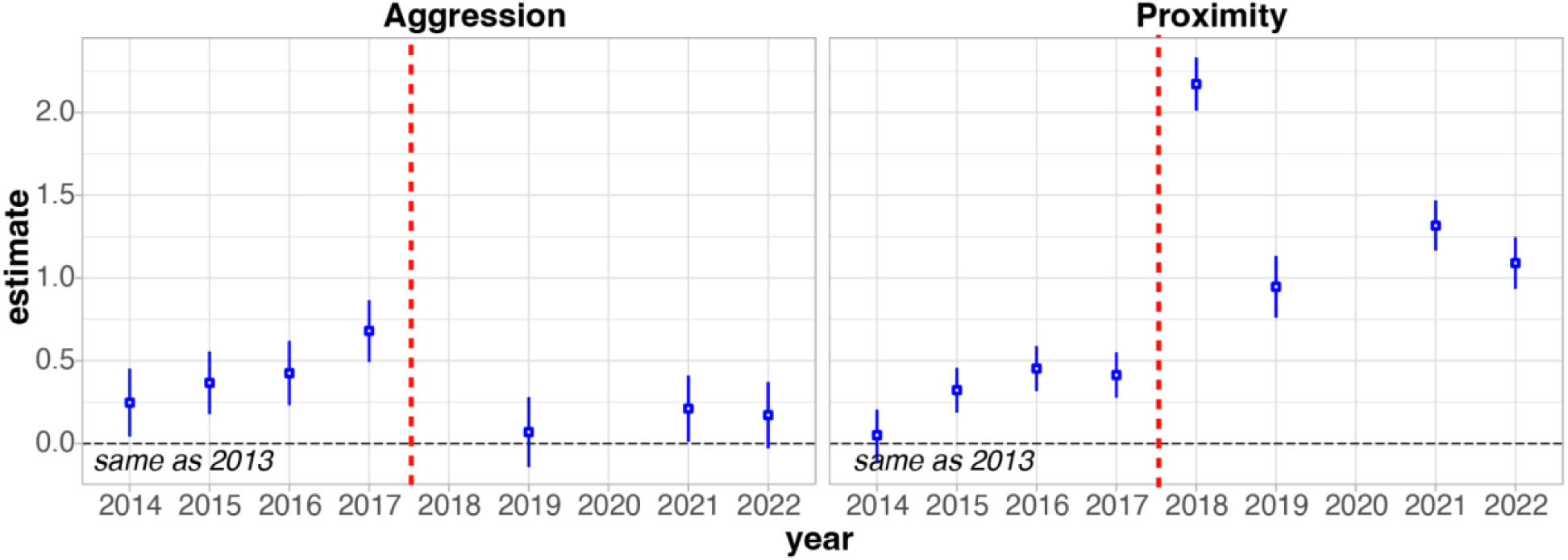
Aggression and proximity in years 2014-2022 relative to 2013 (baseline). The year 2013 was a particularly low aggression year, but most other years pre-hurricane have higher aggression than post-hurricane. See **Data S1** for full model output. Red dotted line: hurricane.

**Fig. S4.**
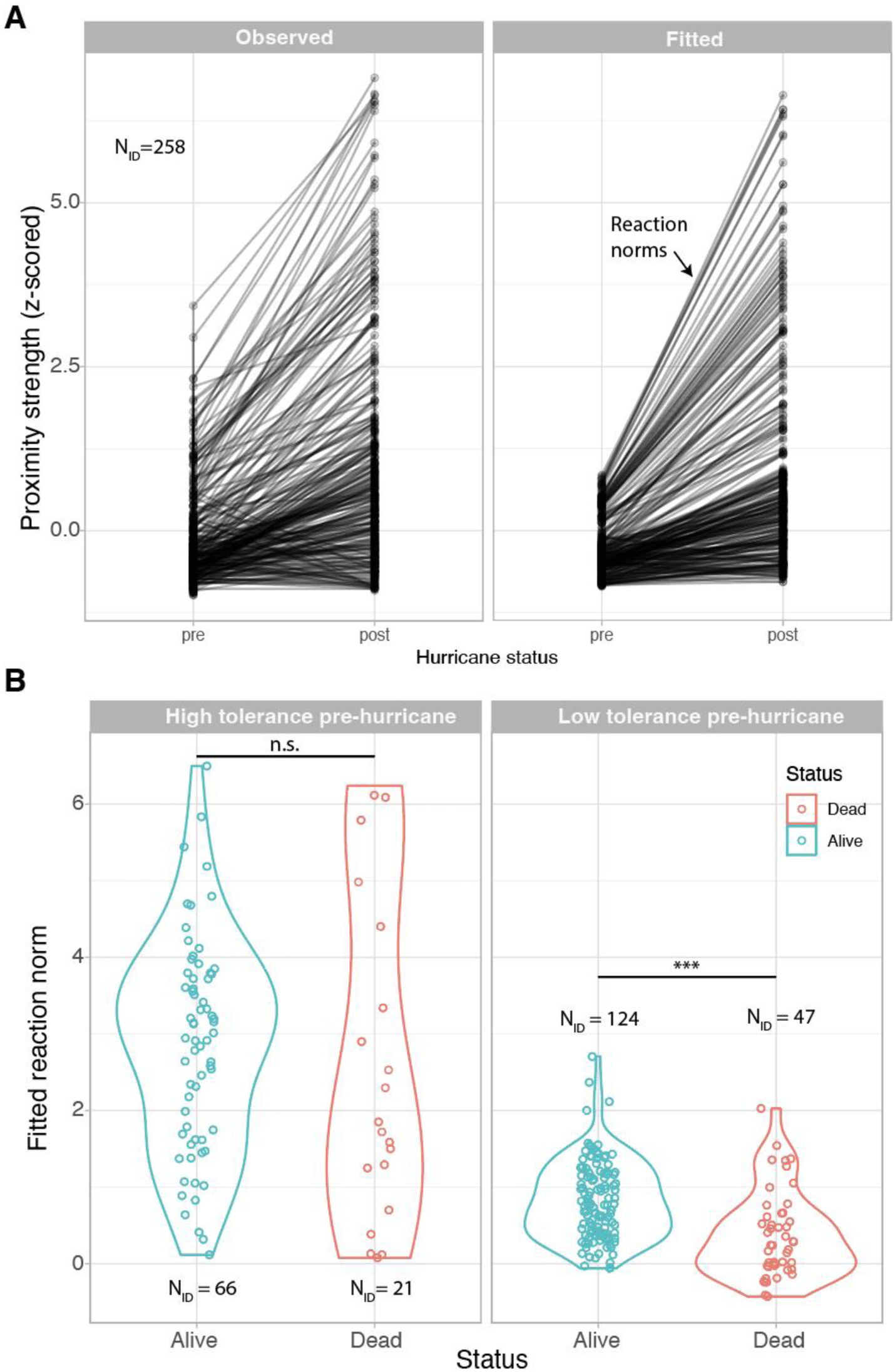
Social plasticity as measured by reaction norms and its relationship with survival. **A**) Observed (left) and fitted (right) plasticity in proximity. Fitted individual slopes from random regression models correspond to reaction norms (see **Suppl. Text**). **B**) Individual variation in reaction norms predict survival for individuals with low tolerance levels pre-hurricane, but not for individuals which were already highly tolerant before the disaster (Bayesian bivariate mixed-effect model, see Suppl. Text). ***: p<0.01; n.s.: not significant.

**Table S1.** Sample sizes by group and year. Note that individuals were sampled on average in 2.7 years.

|  | 2013 | 2014 | 2015 | 2016 | 2017 | 2018 |
| --- | --- | --- | --- | --- | --- | --- |
| **F** | 100 | 106 | 103 | 123 | 113 | 0 |
| **HH** | 0 | 45 | 0 | 69 | 0 | 0 |
| **KK** | 30 | 0 | 54 | 0 | 61 | 63 |
| **R** | 0 | 0 | 94 | 125 | 0 | 0 |
| **S** | 0 | 0 | 0 | 0 | 0 | 0 |
| **TT** | 0 | 0 | 0 | 0 | 0 | 0 |
| **V** | 0 | 0 | 80 | 90 | 102 | 67 |

**Table S2.** Sample size of individuals only pre-hurricane, only post-hurricane and in both periods.

|  |  |
| --- | --- |
| Number of IDs pre-hurricane only | 359 |
| Number of IDs post-hurricane only | 173 |
| Number of IDs both pre- and post-hurricane | 258 |

